# Ca^2+^ binding to Esyt is required to modulate membrane contact site density in *Drosophila* photoreceptors

**DOI:** 10.1101/2024.04.05.588361

**Authors:** Vaisaly R Nath, Harini Krishnan, Shirish Mishra, Padinjat Raghu

## Abstract

Membrane Contact Sites (MCS) between the plasma membrane (PM) and endoplasmic reticulum (ER) have been shown to regulate Ca^2+^ influx into animal cells. However, the mechanisms by which cells modulate ER-PM MCS density is not understood and the role of Ca^2+^, if any, in regulating this process is not known. We report that in *Drosophila* photoreceptors, MCS density is dependent on the activity of the Ca^2+^ permeable channels-TRP and TRPL. This regulation of MCS density by Ca^2+^ is mediated by extended synaptotagmin (dEsyt), a protein localised to ER-PM MCS in photoreceptors and previously shown to regulate MCS density. We find that the Ca^2+^ binding activity of dEsyt is required for its functional activity *in vivo*. dEsyt^CaBM^, a Ca^2+^ non-binding mutant of dEsyt is unable to modulate MCS structure in a manner equivalent to its wild type counterpart. Further, when reconstituted in dEsyt null photoreceptors, in contrast to wild type dEsyt, dEsyt^CaBM^ is unable to rescue ER-PM MCS density and other key phenotypes. Finally, when expressed in wild type photoreceptors, dEsyt^CaBM^ phenocopies loss of dEsyt function. Taken together, our data supports a role for the Ca^2+^ binding activity of dEsyt in regulating the ER-PM MCS density in photoreceptors thus tuning signal transduction in response to Ca^2+^ influx triggered by ambient illumination.

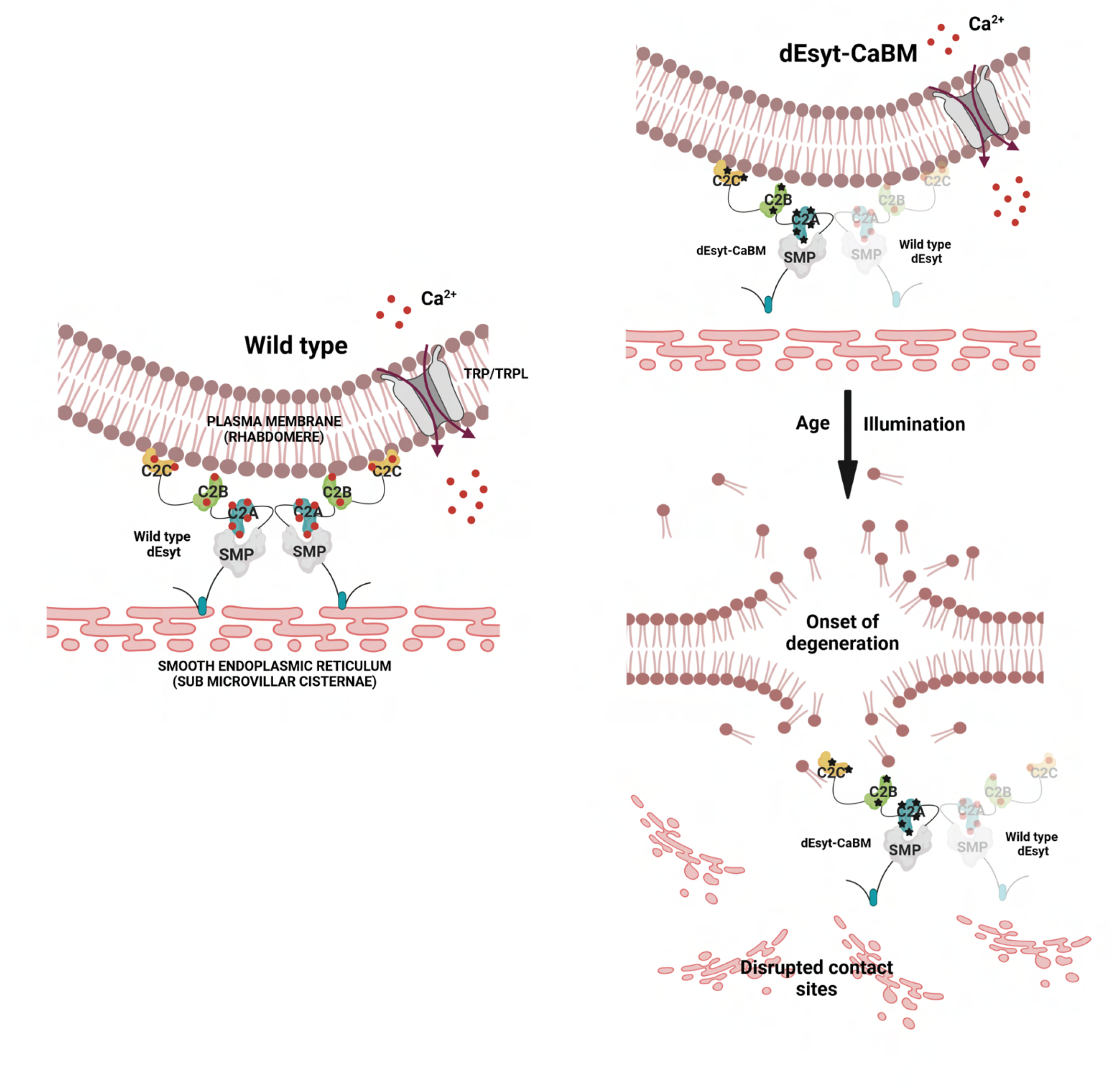

## Introduction

In many eukaryotic cells, the response to stimuli is transduced to cellular responses by changes in intracellular calcium (Ca^2+^) levels (Berridge, 1997). For example, ligand binding to numerous cell surface receptors triggers the activation of phospholipase C (PLC) leading to both Ca^2+^ release from intracellular stores as well as influx from the extracellular compartment into the cytoplasm thus regulating cell physiology (Berridge, 2009). Likewise, in the nervous system, action potentials that mediate neuronal excitation leads to Ca^2+^ influx into the presynaptic terminal through voltage gated Ca^2+^ channels leading to vesicle fusion and neurotransmitter release (Catterall et al., 2013; Dunlap et al., 1995; Mochida et al., 1998; Sabatini and Regehr, 1996; Südhof, 1995). During such events, intracellular Ca^2+^ levels ([Ca^2+^]i) near the plasma membrane (PM) rise and impact the function of proteins at this location. For example, in the context of neurotransmission, the activity of proteins such as synaptotagmin and Munc13 that participate in vesicle fusion and neurotransmitter release is modulated by [Ca^2+^]i (Silva et al., 2021).

In recent years, Membrane Contact Sites (MCS) between the PM and ER have been described as sites for localised biochemical exchange between these organelle membranes (Saheki and De Camilli, 2017). Ca^2+^ signalling and MCS function have been functionally linked. In cells, depletion of intracellular Ca^2+^ reserves increase the areas of apposition between the PM and the ER (Chang et al., 2013; Giordano et al., 2013; Petersen et al., 2017), which supports crucial cellular processes like excitation-contraction coupling in muscles (Endo, 2009). Further, Ca^2+^ influx into cells has been linked to MCS function; the interaction of the ER protein, stromal interaction molecule 1 (STIM1), with the PM Ca^2+^ channel Orai1 leads to its gating thus triggering store operated calcium entry (SOCE) (Prakriya and Lewis, 2015). Several classes of proteins that are localised to ER-PM MCS have been identified. Such proteins, localised within nanoscale distances of the PM are well positioned to detect changes in [Ca^2+^]_i_ at this location and may therefore serve as sensors of cell physiology mediated by Ca^2+^ influx across the PM. One such protein is extended synaptotagmins (Esyt) (Reinisch and De Camilli, 2016). Esyts are ER resident proteins that connect the ER to the PM via their C2 domains. Some of the C2 domains of Esyt have been reported to be regulated by Ca^2+^; for example, in Esyt1 it is reported that extracellular Ca^2+^ influx enhance ER-PM MCS through binding to the C2 domains (Idevall-Hagren et al., 2015) followed by a Ca^2+^ dependent release of an interdomain autoinhibitory mechanism has been proposed (Bian et al., 2018). In mammalian cells, it has also been proposed that Ca^2+^ binding to the C2 domains may regulate the lipid transfer function of the synaptotagmin like mitochondrial lipid binding protein (SMP) domain (Bian et al., 2018; Idevall-Hagren et al., 2015) suggesting the Ca^2+^ regulation of Esyt function during signalling. While these findings indicate that the C2 domains of Esyt can bind and respond to Ca^2+^, the relevance of this Ca^2+^ binding to the physiological functions of Esyt *in vivo* remains unknown.

In *Drosophila* photoreceptors, transduction of photon absorption is mediated by G-protein coupled PLC signalling leading to a large rise in [Ca^2+^]_i_ levels (Hardie and Raghu, 2001). This rise in [Ca^2+^]_i_ is mediated by the opening of the light sensitive, Ca^2+^ permeable PM channels: transient receptor potential (TRP) and *trp*-like (TRPL) (Reuss et al., 1997). The apical plasma membrane in *Drosophila* photoreceptors is expanded to form finger like microvilli and PLC activation and Ca^2+^ influx through TRP and TRPL channels occurs at the microvillar PM. At the base of the microvilli, the ER is closely arranged next to the microvillar PM to form an ER-PM MCS **(Figure 1A)**. During PLC signalling, in the context of phototransduction, lipid transfer occurs at this MCS through the PI and PA transfer activity of the RDGB protein (Yadav et al., 2015)[reviewed in (Raghu et al., 2021)] and the *Drosophila* photoreceptor has been an influential model system for the understanding of ER-PM MCS biology *in vivo* (Yadav et al., 2016). The *Drosophila* genome contains a single gene encoding Esyt (*dEsyt*) (Kikuma et al., 2017; Nath et al., 2020) and loss of function mutants of *dEsyt* leads to loss of ER-PM integrity, mislocalisation of RDGB and consequent retinal degeneration (Nath et al., 2020). In *Drosophila* photoreceptors, dEsyt is localised to the base of the microvillus at the ER-PM MCS and is therefore well positioned to sense and respond to changes in [Ca^2+^]_i_ arising due to activation of TRP and TRPL during phototransduction. However, the regulation of ER-PM MCS and dEsyt by [Ca^2+^]_i_ is not known. Here, we report that TRP and TRPL channel function is required to maintain normal MCS density in *Drosophila* photoreceptors. Using a Ca^2+^ non-binding mutant of dEsyt (dEsyt^CaBM^), we find that loss of Ca^2+^ binding activity leads to both mislocalisation of the protein and the mutant protein phenocopies loss of dEsyt function. Thus, dEsyt binds Ca^2+^ and mediates the transduction of photoreceptor illumination into MCS density *in vivo*. This in turn tunes the photoreceptor function to ongoing ambient illumination.

**Figure 1:**
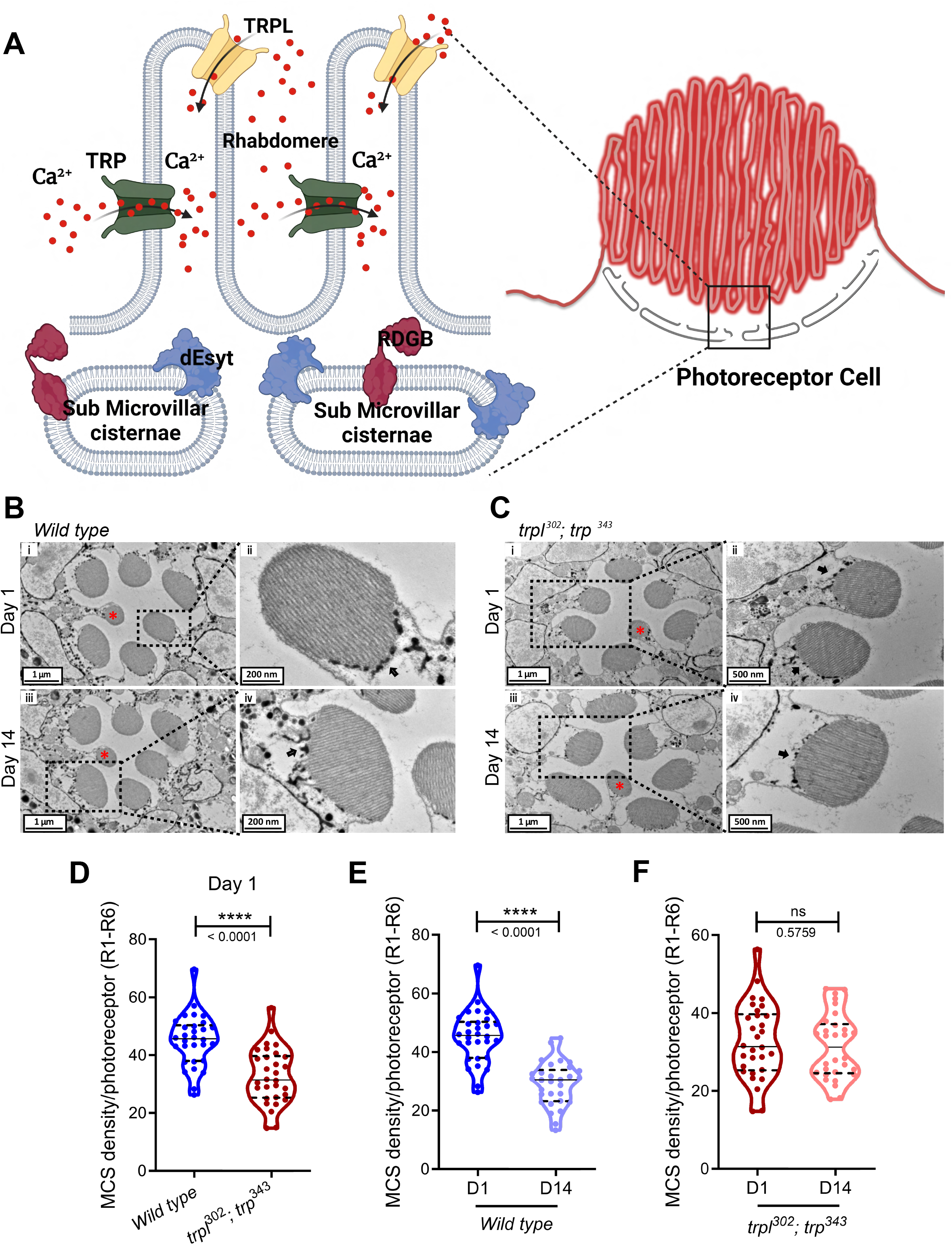
MCS density is regulated by TRP channels in *Drosophila* photoreceptors. **(A)** Schematic showing the spatial orientation and dimensions of the location of TRP, TRPL, dEsyt and RDGB across the microvillar plasma membrane and the sub-microvillar cisternae in *Drosophila* photoreceptor cell. **(B)** TEM images of a single ommatidium from *Wild type* photoreceptors of flies reared in dark **(i)** day 1 (D1) and **(iii)** day 14 (D14), scale bar: 1µM. (* - R7 photoreceptor). Magnified image showing a single photoreceptor from the same ommatidium **(ii)** day1 and **(iv)** day 14. Scale bar: 200 nm. Arrow indicates the SMC forming an MCS with the microvillar PM. **(C)** TEM images of a single ommatidium from *trpl^302^;trp^343^* photoreceptors of flies reared in dark **(i)** day 1 (D1) and **(iii)** day 14 (D14), scale bar: 1µM. (* - R7 photoreceptor). Magnified image showing a single photoreceptor from the same ommatidium **(ii)** day1 **(iv)** day 14. Scale bar: 500 nm. Arrow indicates the SMC forming an MCS with the microvillar PM. **(D)** Quantification of the number of MCS per photoreceptor of *Wild type* day 1 v/s *trpl^302^;trp^343^*day 1 old flies reared in dark, X-axis indicates the genotype and Y-axis indicates the number of MCS/photoreceptor. n=30 photoreceptors from 3 separate flies (R1-R6) [N=3; n=30]. **(E)** Quantification of the number of MCS per *Wild type* photoreceptor of day 1 and day 14 old flies reared in dark (from figure 1B), X-axis indicates the genotype and age of the fly. Y-axis indicates the number of MCS/photoreceptor. n=30 photoreceptors from 3 separate flies (R1-R6) [N=3; n=30]. **(F)** Quantification of the number of MCS per *trpl^302^;trp^343^* photoreceptor of day 1 and day 14 old flies reared in dark (from figure 1C), Y-axis indicates the number of MCS/photoreceptor. n=30 photoreceptors from 3 separate flies (R1-R6) [N=3; n=30]. *Violin plots with mean ± SD are shown. Statistical tests: (D, E and F) Student’s unpaired t test. ns - Not significant; ****p value <0.0001*.

## Results

### TRP channels regulate MCS density in *Drosophila* photoreceptors

Age-related decrease in ER-PM MCS density in wild-type photoreceptors are reliant on continued PLC activity (Nath et al., 2020). A key outcome of light induced PLC activity in photoreceptors is the activation of the light-sensitive calcium- and cation-selective plasma membrane channels encoded by *trp* and *trpl* **(Figure 1A)** (Hardie and Minke, 1992; Reuss et al., 1997). To test the effect of calcium influx, if any on MCS density, we measured the density of ER-PM MCS in *trpl^302^; trp^343^* mutants compared to controls **(Figure 1B, C)**. At eclosion, MCS density in *trpl^302^; trp^343^* was approximately 30% compared to 45% in controls **(Figure 1D)**. As previously reported (Nath et al., 2020), wild type flies show an age dependent drop in MCS density **(Figure 1E)**. However, unlike wild-type flies, *trpl^302^; trp^343^* photoreceptors did not exhibit any age-related drop in MCS density; hence, 14-day-old *trpl^302^; trp^343^* photoreceptors displayed the same MCS density as 1-day-old flies **(Figure 1F)**. Therefore, MCS density in aging photoreceptors is dependent on TRP and TRPL channel activity.

### Calcium binding residues of dEsyt C2A domain are structurally similar to hEsyt2 C2A domain

The tethering and lipid transfer activity of mammalian Esyts are reported to be influenced by Ca^2+^ (Giordano et al., 2013; Saheki et al., 2016; Yu et al., 2016). However, the sensitivity of dEsyt to Ca^2+^ binding is not known. C2 domains from several Esyt proteins were aligned using clustalO (Sievers et al., 2011) to confirm the conservation of calcium binding residues **(Figure 2A)**. *dEsyt* contains nine conserved calcium binding residues spread across the three C2 domains-C2A (D364, D374, D421, D423, E429); C2B (D517, D564) and C2C (D746, D752) (Kikuma et al., 2017). The C2A and C2B domains from human synaptotagmin, C2A-C2E domains from human Esyt1 (*hEsyt1*), C2A-C2C domains from human Esyt2 (*hEsyt2*) and C2A-C2C domains of dEsyt were aligned. The Ca^2+^ binding residues of the dEsyt C2A domain were conserved and were similar to the residues in hEsyt2 **(Figure 2A)**. The X-ray crystallographic structure of hEsyt2 C2A and C2B domain (PDB ID: 4npk) bound to Ca^2+^ shows that the C2A domain can bind calcium while the C2B domain does not bind calcium (Xu et al., 2014). The binding of the first calcium ion to C2A domain occurs with high affinity (K_d_ < 10µM) while the binding of other two calcium ions has lower affinity (K_d_ = 100-200 µM) (Xu et al., 2014). The sequence identity between hEsyt2 and dEsyt protein sequences is 49.52% **(Figure 2B)**. We generated the structure of dEsyt C2A domain by *in-silico* homology modelling using MODELLER (Webb and Sali, 2014) with the hEsyt2 structure as template (PDB ID: 4npk). The overall RMSD between the two structures is 0.159 Å **(Figure 2C)** while the RMSD of the 5 residues that interact with Ca^2+^ is less than 0.1 Å **(Figure 2E)**. This finding suggests that the interaction of Ca^2+^ binding residues of dEsyt is similar to hEsyt2. To check which residues are capable of forming Ca^2+^ co-ordination in the dEsyt structure, we used the MIB, a metal ion binding site prediction tool (Lin et al., 2016). The residues D364, D374, D421, D423 and E429 of dEsyt protein were predicted with scores greater than 3 and marked as residues important for Ca^2+^ binding **(Figure 2D)**. These residues correspond to the five aspartate residues of hEsyt2 as shown in the PDB structure of 4npk. The four aspartate residues were then mutated to asparagine and the glutamate residue was mutated to glutamine using the “build model” module of FOLDX (Schymkowitz et al., 2005). The MIB tool was used to check residues interacting with calcium ions in the mutant dEsyt structure. All the five residues in the wild type protein structure that were predicted to bind calcium now showed lower MIB score and reduced ion binding capacity **(Figure 2F)**. This finding strongly suggests that these aspartate and glutamate residues play an important role in interaction with Ca^2+^.

**Figure 2:**
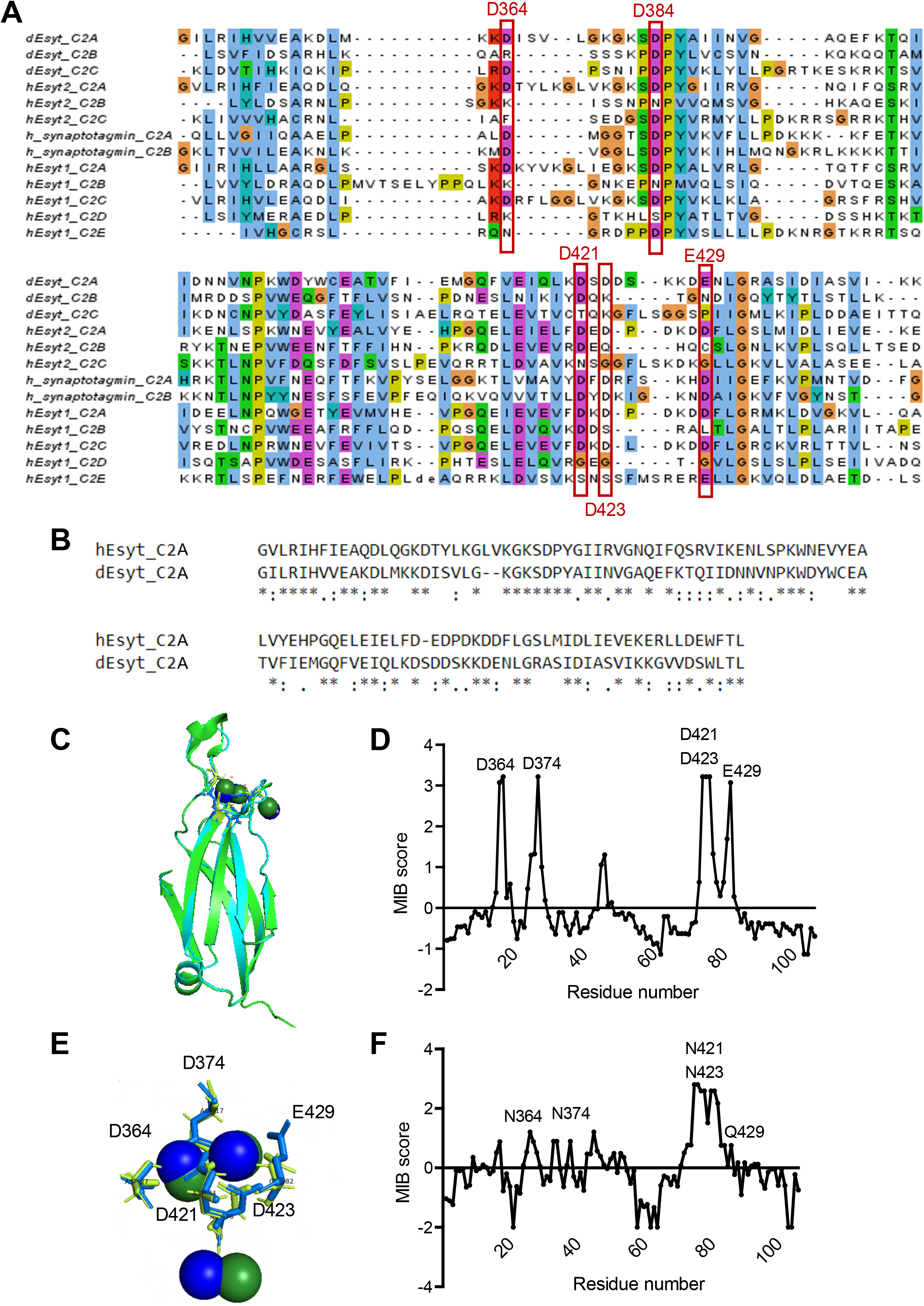
Calcium binding residues of dEsyt C2A domain are structurally similar to hEsyt2 C2A domain. **(A)** Alignment of C2 domain from dEsyt (C2A-C), hEsyt2 (C2A-C), h-Synaptotagmin (C2A-B) and hEsyt1 (C2A-E). The residues important for binding calcium ions have been marked in black rectangles. **(B)** Alignment between hEsyt2 and dEsyt C2A domain used for homology modelling using MODELLER. **(C)** Structural superposition of template hEsyt2 PDB ID 4npk and dEsyt model generated by MODELLER (RMSD=0.159 Å). **(D)** MIB scores for residues of the wild type dEsyt protein. **(E)** Structural superposition of residues involved in calcium metal coordination. **(F)** MIB scores for residues of the mutant dEsyt protein.

### Calcium binding to dEsyt is required for localisation to ER-PM contact sites

To test the sensitivity of dEsyt to Ca^2+^ binding and its impact on cell physiology *in vivo,* we generated a mutant version of dEsyt that cannot bind Ca^2+^. We generated a dEsyt transgene where the four conserved aspartates (D) were mutated to asparagine (N) and the glutamic acid (E) to glutamine (Q) in the C2A of dEsyt. Further, each of the two conserved aspartates (D) in C2B and C2C were mutated to asparagine (N). This transgene carrying a total of nine mutations should render the protein unable to bind calcium (*dEsyt^CaBM^*) **(Figure 3A)**. Aspartate-to-asparagine mutations (D→N) result in inhibition of Ca^2+^ binding by removing the negative charge from the calcium binding pocket and result in minimal disruption of the protein structure as the mutations partially mimic Ca^2+^ binding (Fernández-Chacón et al., 2002; Robinson et al., 2002). We tested the requirement of intact Ca^2+^ binding residues in dEsyt for its localisation. On expression in *Drosophila* S2R+ cells, dEsyt^CaBM^::GFP was found to primarily localise to the PM with a fraction of the protein still residing in certain membrane compartments within the cytosol **(Figure 3B)**. This is in contrast to wild type dEsyt::GFP that is primarily localised in the membrane compartments within the cytosol. We also studied the localisation of dEsyt^CaBM^::GFP in *Drosophila* photoreceptors. In photoreceptors, *Rh1>dEsyt::GFP* appears localised towards the base of the rhabdomere; by contrast, *Rh1>dEsyt^CaBM^::GFP* was found to be exclusively localised to the rhabdomeres (apical plasma membrane) **(Figure 3C)**. Thus, the Ca^2+^ binding activity of dEsyt is important to retain the protein at the ER-PM contact sites and in the absence of Ca^2+^ binding, the protein appears mainly localised to the PM.

**Figure 3:**
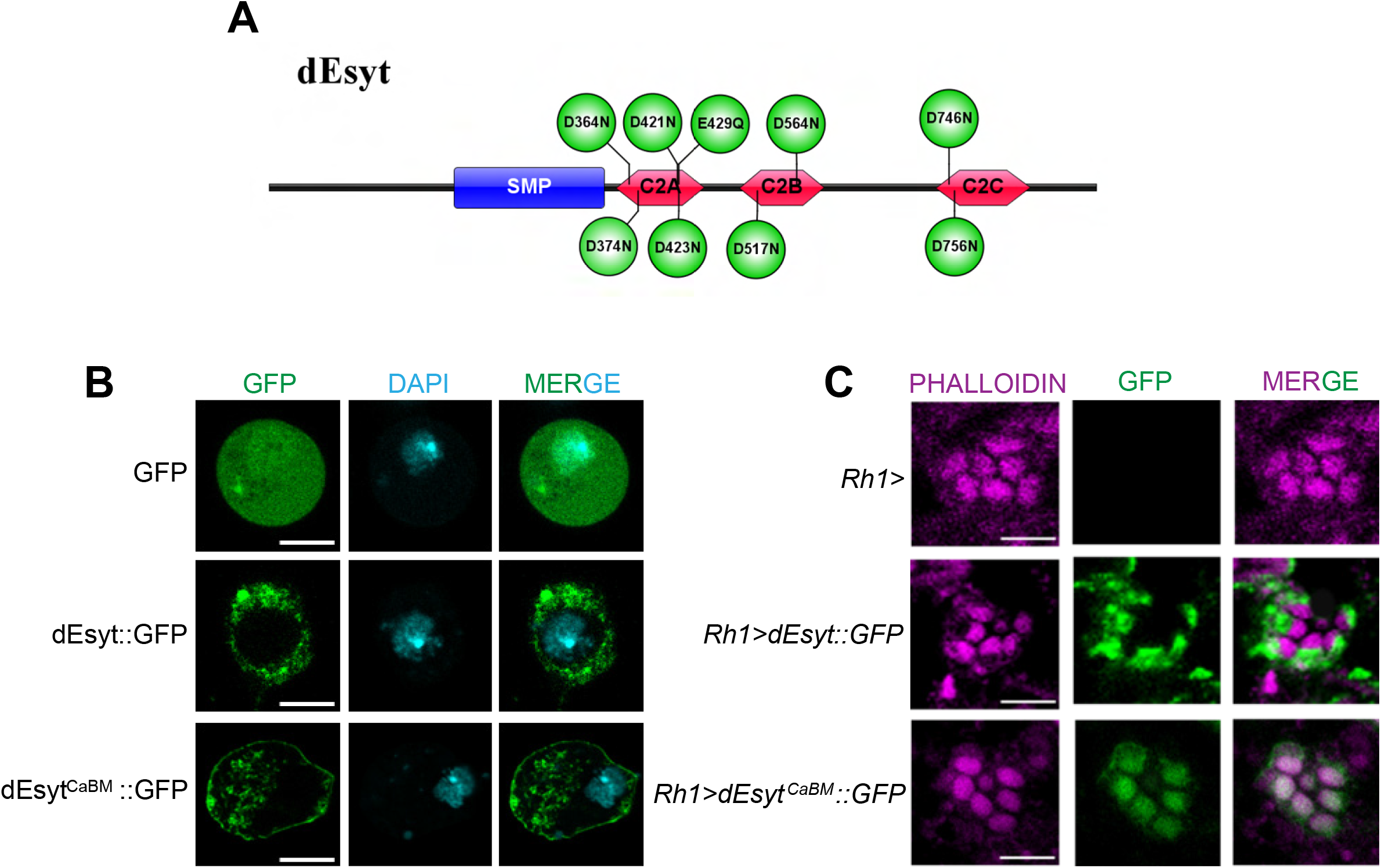
Localisation of dEsyt to the ER-PM contact sites require calcium binding to dEsyt. **(A)** Schematic showing the site directed mutations introduced into the nine conserved calcium binding residues spread across the three C2 domains of dEsyt. **(B)** Confocal images of S2R+ cells transfected with pUASt-attB::GFP, pUASt-attB-dEsyt::GFP and pUASt-attB-dEsyt^CaBM^::GFP. Green represents signal from GFP. DAPI (cyan) stains the nucleus. **(C)** Confocal images showing the localisation of exogenously expressed dEsyt::GFP and dEsyt^CaBM^::GFP driven using eye specific Rh1-Gal4 in 1-day old dark reared flies. Rh1-Gal4 is shown as a control. A single ommatidium is shown. Scale bar: 5 µM. Phalloidin marks F-actin staining and highlights rhabdomeres R1-R7.

### Functional activity of *dEsyt^CaBM^*

To test the function of dEsyt^CaBM^ in cells, we overexpressed this protein in *Drosophila* photoreceptors. When dEsyt::GFP is expressed in wild type photoreceptors (*Rh1> dEsyt::GFP*), it does not affect photoreceptor structure even when flies are grown in 5000 Lux constant illumination **(Figure 4A,B)**. However, when dEsyt^CaBM^::GFP is expressed under equivalent conditions (*Rh1> dEsyt^CaBM^::GFP*), this resulted in a light dependent retinal degeneration **(Figure 4A, B)**. The kinetics of this degeneration closely paralleled with that previously reported for *dEsyt^KO^* (Nath et al., 2020) suggesting that dEsyt^CaBM^::GFP might exert a dominant negative effect on the endogenous protein.

**Figure 4:**
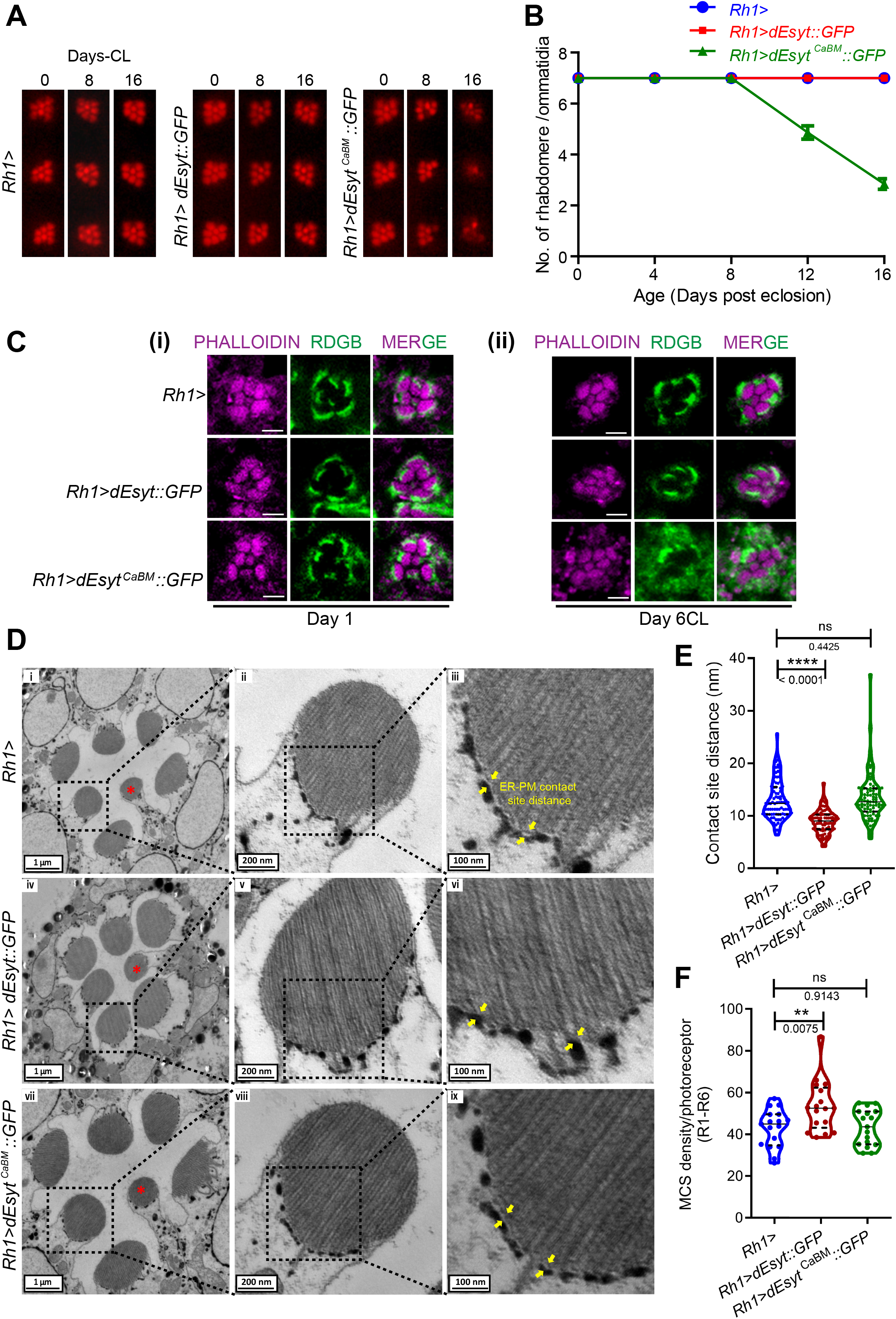
Functional activity of *dEsyt^CaBM^*. **(A)** Representative optical neutralization (ON) images showing rhabdomere structure of the indicated genotypes. Rearing conditions and the age of the flies are indicated on top. (*CL-Constant Light). **(B)** Quantification of the time course taken for retinal degeneration. 50 ommatidia were scored from 5 flies of each genotype and plotted. [N=5; n=50] **(C)** Confocal images showing the localisation of RDGB in *Rh1>*, *Rh1>dEsyt::GFP* and *Rh1>dEsyt^CaBM^::GFP* photoreceptors of flies which are (i) 1- day old dark reared (ii) 6 days old-exposed to constant illumination (5000 Lux) For (i) and (ii) RDGB visualized is detected using an antibody against the endogenous protein. Rhabdomeres are outlined using phalloidin which marks F-actin. A single ommatidium is shown. Scale bar: 5 µM. **(D)** TEM images of a single ommatidium from photoreceptors of flies reared in dark, 1-day-old (i) *Rh1>*, (iv) *Rh1>dEsyt::GFP* and (vii) *Rh1>dEsyt^CaBM^::GFP.* Scale bar: 1µM. (* - R7 photoreceptor). (ii, v, viii) Magnified image showing a single photoreceptor from the ommatidium image shown on the left. Scale bar: 200 nm. (iii, vi, ix) Magnified image showing the MCS distance (yellow arrows highlight the distance between ER and PM) of the photoreceptor image shown on the left. Scale bar: 50 nm. **(E)** Quantification of the contact site distance between the apical PM and the associated SMC, X-axis indicates the genotypes and Y-axis indicates the contact site distance in nm, 75 data points from 25 photoreceptors from 3 separate flies (R1-R6). [N=3; n=25] **(F)** Quantification of the MCS density per photoreceptor, X-axis indicates the genotype and Y-axis indicates the MCS density/photoreceptor, n=25 photoreceptors from 3 separate flies (R1-R6). [N=3; n=25] *(B) XY plots with mean ± SD are shown. (E and F) Violin plots with mean ± SD are shown. Statistical tests: (E and F) Student’s unpaired t test. ns - Not significant; **p value<0.01; ****p value <0.0001*.

To understand if altered localisation of dEsyt^CaBM^ in a different compartment (apical PM) within these highly polarised cells can manifest an effect on the ER-PM integrity and thereby contribute to photoreceptor degeneration, we assessed the localisation of a known contact site protein, RDGB, in *Rh1>dEsyt^CaBM^::GFP* lines. At day-1 post eclosion, just like in the control photoreceptors, RDGB was solely localised to the MCS in both *Rh1>dEsyt::GFP* and *Rh1>dEsyt^CaBM^::GFP* **(Figure 4C i)**. However, after six days of continuous illumination, despite the presence of wild type gene, *Rh1>dEsyt^CaBM^::GFP* exhibited diffused localisation of RDGB throughout the photoreceptor cell body **(Fig 4C ii)**; this phenocopies that seen in *dEsyt^KO^* photoreceptors (Nath et al., 2020). In contrast, in wild-type and *Rh1>dEsyt::GFP* photoreceptors, RDGB localisation to the MCS was preserved under identical conditions **(Fig 4C ii)**.

Contact site disruption has previously been shown to be the cause of RDGB mislocalisation in *dEsyt^KO^* photoreceptors (Nath et al., 2020). To determine if the mislocalisation of RDGB in dEsyt^CaBM^ is as a result of a change in contact site density or the distance between the ER and PM, we analysed the photoreceptor ultrastructure. The distance between the ER and PM in wild type photoreceptors is ∼12 - 13 nm (Krishnan et al., 2023) **(Figure 4D i, ii, iii, 4E)**. This contact site gap was substantially less at day-1 post-eclosion in *Rh1>dEsyt::GFP*, showing the protein’s tethering role in bringing the two membranes together. By contrast, the Ca^2+^ binding mutant was unable to reduce contact site distance **(Figure 4D iv-ix, 4E)**. When the same experimental set was examined for MCS density, we discovered that the density enhanced by 10% in *Rh1>dEsyt::GFP* while being comparable between wild type and *Rh1> dEsyt^CaBM^::GFP* flies. **(Figure 4D i-ix, 4F)**.

### dEsyt^CaBM^ can partially rescue the phenotypes of *dEsyt^KO^*

We reconstituted dEsyt^CaBM^::GFP in photoreceptors of *dEsyt^KO^*, a protein null allele (Nath et al., 2020). As a test of function, we monitored the retinal degeneration phenotype of *dEsyt^KO^*. When grown in illumination, *dEsyt^KO^* photoreceptors undergo light dependent retinal degeneration starting at day-8 post-eclosion; reconstitution with dEsyt::GFP (*Rh1> dEsyt::GFP; dEsyt^KO^*) was able to fully rescue this phenotype **(Fig 5 A, B)**. By contrast, reconstitution with dEsyt^CaBM^::GFP (*Rh1> dEsyt^CaBM^::GFP; dEsyt^KO^*) was not able to rescue the retinal degeneration phenotype of *dEsyt^KO^* although it modestly delayed the onset of degeneration from 8 days post-eclosion in *dEsyt^KO^* to 12 days in *Rh1> dEsyt^CaBM^::GFP; dEsyt^KO^***(Fig 5A,B)**. Consistent with this observation, when exposed to constant illumination for 6 days post-eclosion, the localisation of RDGB was rescued in *Rh1>dEsyt::GFP; dEsyt^KO^* and *Rh1>dEsyt^CaBM^::GFP; dEsyt^KO^* **(Figure 5C ii)** by which point RDGB was mislocalised in *dEsyt^KO^* and *dEsyt^CaBM^* photoreceptors.

**Figure 5:**
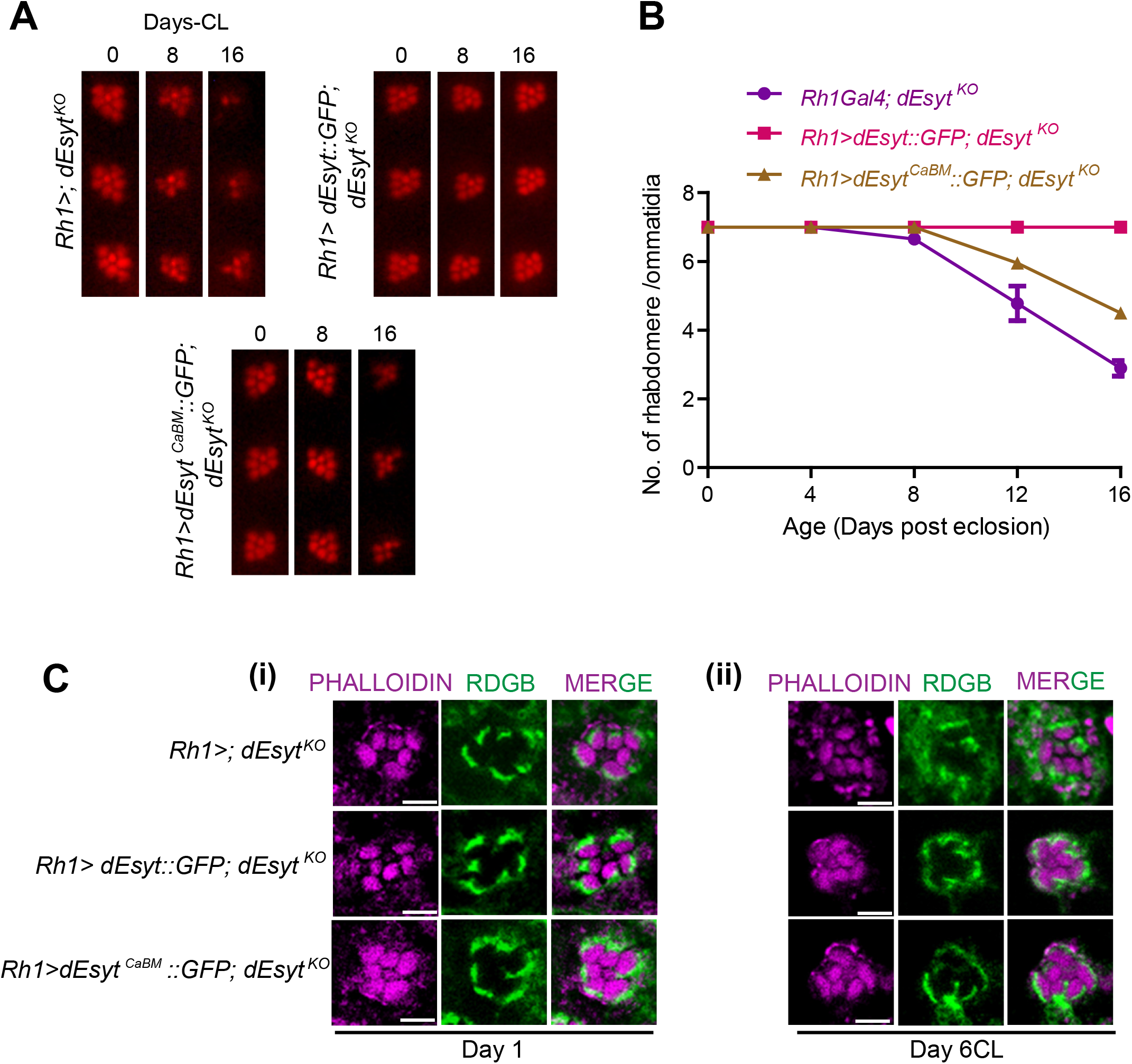
Partial rescue of *dEsyt^KO^* phenotypes by dEsyt^CaBM^. **(A)** Representative optical neutralization (ON) images showing rhabdomere structure of the indicated genotypes. Rearing conditions and the age of the flies are indicated on top. (*CL-Constant Light). **(B)** Quantification of the time course taken for retinal degeneration. 50 ommatidia were scored from 5 flies of each genotype and plotted. [N=5; n=50] **(C)** Confocal images showing the localisation of RDGB in *Rh1>; dEsyt^KO^*, *Rh1>dEsyt::GFP; dEsyt^KO^* and *Rh1>dEsyt^CaBM^::GFP; dEsyt^KO^* photoreceptors of flies which are (iii) 1- day old dark reared (iv) 6 days old- exposed to constant illumination (5000 Lux) For (i) and (ii) RDGB visualized is detected using an antibody against the endogenous protein. Rhabdomeres are outlined using phalloidin which marks F-actin. A single ommatidium is shown. Scale bar: 5 µM.

We examined the MCS density in *dEsyt^KO^* photoreceptors reconstituted with dEsyt::GFP and dEsyt-CaBM::GFP. Compared to *dEsyt^KO^* **(Figure 6A i-ii, D)**, for R1-R6 photoreceptors, MCS density was marginally higher in *Rh1>dEsyt::GFP; dEsyt^KO^***(Figure 6B i-ii, D)** and *Rh1>dEsyt^CaBM^::GFP; dEsyt^KO^* **(Figure 6C i-ii, D)** at eclosion indicating that altering the calcium-binding residues in dEsyt does not affect the formation of ER-PM contact sites during the *Drosophila* photoreceptor development. We then measured MCS density in *Rh1>dEsyt^CaBM^::GFP; dEsyt^KO^* as a function of age. While ER-PM MCS was nearly non-existent at Day 14 in *dEsyt^KO^* (Nath et al., 2020) **(Figure 6A iii, iv; E)**, *Rh1>dEsyt::GFP; dEsyt^KO^* showed complete phenotypic rescue and the number of MCS was comparable to Day 1 **(Figure 6B i-iv; D, E)**. As a function of age, from Day 1 to Day14, the MCS density was reduced by 20% in *Rh1>dEsyt^CaBM^::GFP; dEsyt^KO^* **(Figure 6C iii, iv; E)**. Thus, the calcium binding mutant of the protein was able to partially rescue the contact site integrity of *dEsyt^KO^* at Day 14 **(Figure 6E)**.

**Figure 6:**
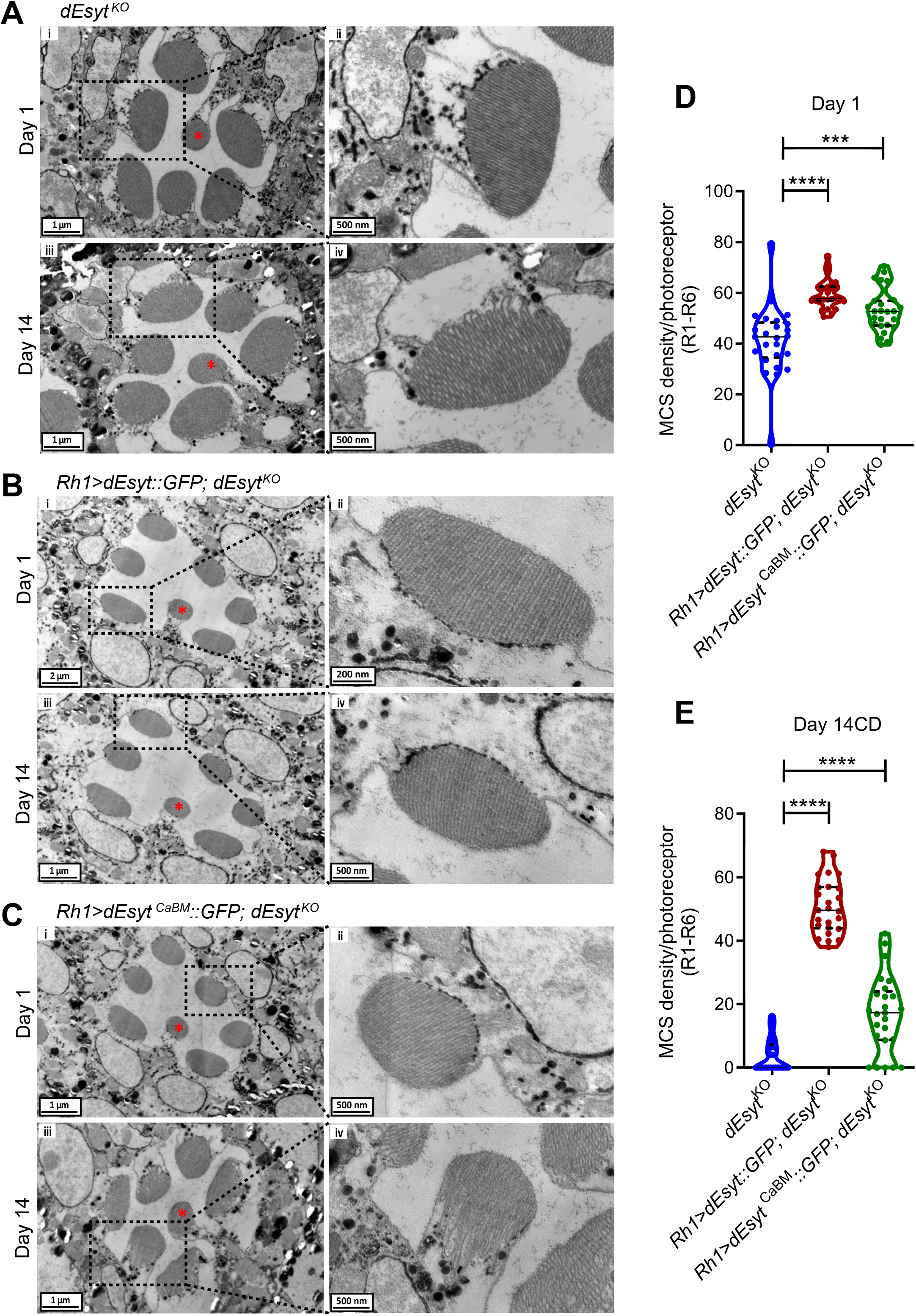
Partial rescue of *dEsyt^KO^* phenotypes by dEsyt^CaBM^. TEM images of a single ommatidium from photoreceptors of flies reared in dark, scale bar: 1 µM. (* - R7 photoreceptor). **(A) (i)** *dEsyt^KO^*- day 1 **(iii)** *dEsyt^KO^*- day 14 **(B) (i)** *Rh1>dEsyt::GFP; dEsyt^KO^*- day 1 **(iii)** *Rh1>dEsyt::GFP; dEsyt^KO^* - day 14 **(C) (i)** *Rh1>dEsyt^CaBM^::GFP; dEsyt^KO^*- day1 **(iii)** *Rh1>dEsyt^CaBM^::GFP; dEsyt^KO^* - day 14 (Ai, iii, Biii, Ci, iii) Scale bar: 1um; (Bi) Scale bar: 2um. **(A-ii, iv; B-ii, iv; C-ii, iv)** Magnified image showing a single photoreceptor from the ommatidium image shown on the left. (Bii) Scale bar: 200 nm; (Aii, iv, Biv, Cii, iv) Scale bar: 500 nm. **(D)** Quantification of the number of MCS per photoreceptor of day-1-old dark-reared flies, X-axis indicates the genotype and Y-axis indicates the number of MCS/photoreceptor, n=25 photoreceptors from 3 separate flies (R1-R6). [N=3; n=25] **(E)** Quantification of the number of MCS per photoreceptor of flies reared in 14 Day CD (Constant Dark), X-axis indicates the genotype and age of the flies and Y-axis indicates the number of MCS/photoreceptor, n=25 photoreceptors from 3 separate flies (R1-R6). [N=3; n=25] *(D and E) Violin plots with mean ± SD are shown. Statistical tests: Student’s unpaired t test. ***p value<0.001; ****p value <0.0001*.

## Discussion

Since ER-PM junctions are located within nanoscale distances of the PM, proteins localised to this site are ideally positioned to detect Ca^2+^ influx across the PM and this information can then be used to mediate sub-cellular responses and tune cellular physiology. For e.g during sensory transduction such as that which occurs in photoreceptors, detection of Ca^2+^ influx, the final outcome of phototransduction at ER-PM MCS can be used to tune ongoing signalling. In *Drosophila* photoreceptors, sensory transduction depends on MCS function thus providing a setting to link MCS function with sensory transduction. In this scenario, proteins localised to ER-PM MCS that can bind Ca^2+^ are essential to couple sensory transduction with MCS density. Previous studies have shown that in mammalian cells, Esyts that are localised to ER-PM MCS can bind to Ca^2+^ and that the function of the molecule can be regulated by Ca^2+^ binding (Bian et al., 2018; Giordano et al., 2013; Idevall-Hagren et al., 2015; Xu et al., 2014; Yu et al., 2016). However, the relevance of such Ca^2+^ binding to the regulation of physiology was unknown. In this study, using *Drosophila* photoreceptors, a model for ER-PM MCS function *in vivo*, we find that the reported modulation of ER-PM MCS by light and age (Nath et al., 2020) depends on Ca^2+^ influx through TRP channels. PLC activity is required for TRP channel activation in photoreceptors (Raghu et al., 2012). It has previously been reported that intact PLC function is required for modulation of MCS density in *Drosophila* photoreceptors (Nath et al., 2020). Taken together, the observation that both PLC activity and TRP channel function are required for MCS density modulation underscores the role of phototransduction mediated PM Ca^2+^ influx in regulating MCS density *in vivo* and as such also reports the modulation of MCS density by ongoing changes in the environment (in this case light). Thus, in a signal transduction system, our observation establishes a link between MCS density and ongoing signal transduction.

How might Ca^2+^ influx, the output of sensory transduction be transduced into modulation of MCS density? In principle this requires a protein that is localised to the MCS to bind Ca^2+^ and for its activity to be transduced into modulation of MCS density. In photoreceptors, dEsyt is an ER-PM MCS localised protein (Nath et al., 2020) and overexpression of dEsyt in wild type photoreceptors is able to increase MCS density and reduce ER-PM distance at the MCS (this study Fig 4). How might dEsyt transduce ongoing transduction into altered MCS density? Our *in silico* analysis reveals that evolutionarily conserved residues in the C2A domain of dEsyt are required for Ca^2+^ binding. When these residues are mutated, the non- Ca^2+^ binding version of dEsyt (dEsyt^CaBM^) so generated shows reduced function with respect to MCS modulation. Overexpression of dEsyt^CaBM^ led to no change in ER-PM MCS density and ER-PM distance compared to equivalent overexpression of wild type dEsyt (Fig 4E, F). These findings imply that Ca^2+^ binding to dEsyt is required for the ability of the overexpressed protein to modulate MCS function. It has previously been demonstrated that endogenous dEsyt is required to maintain MCS density in photoreceptors (Nath et al., 2020). Is the Ca^2+^ binding activity of endogenous dEsyt required for this function? We found that reconstitution of *dEsyt^KO^* with wild type dEsyt was able to restore the reduced MCS density but dEsyt^CaBM^ was less effective in doing so. This finding underscores the importance of Ca^2+^ binding to dEsyt for mediating its ability to modulate MCS density *in vivo*.

The ability of Esyt to sense Ca^2+^ influx across the PM and modulate MCS density should require the protein to be positioned accurately at ER-PM MCS. Previous studies have described motifs required to correctly position Esyt at the ER-PM contact site, such as an N-terminal ER membrane-associated region and the calcium induced C2 domain-dependent interaction with phosphatidylinositol 4,5-bisphosphate (PIP_2_) to bind the PM (Chang et al., 2013; Giordano et al., 2013). In this study, we found that mutating the calcium binding residues (dEsyt^CaBM^), repositioned dEsyt away from its normal location at ER-PM MCS to the rhabdomeral plasma membrane and this was associated with its inability to function normally, implying that dEsyt^CaBM^ exerts a dominant negative effect on wild type dEsyt. One possible mechanism for the phenotypes exhibited by dEsyt^CaBM^ expression in wild type cells is suggested by the findings of a structural and mass spectrometry investigation of hEsyt2 that reveals that the SMP domain dimerizes to create a 90Å long cylinder to facilitate the transfer of lipids (Schauder et al., 2014). Other mechanisms may also operate but remains to be understood. Consistent with our observations, in a previous study, *dEsyt^D-N^* flies, which lack the negatively charged amino acids in each C2 domain necessary for Ca^2+^ binding, protein expression was restricted to the cell body rather than the presynaptic terminals, and resulted in larval lethality (Kikuma et al., 2017).

It is still unclear whether the three C2 domains have different binding affinities for calcium as previously reported for a mammalian Esyt (Bian et al., 2018) and what their response would be to the high Ca^2+^ concentrations experienced by the region of the ER-PM MCS in *Drosophila* photoreceptors. Nonetheless, this study demonstrates the role of Ca^2+^ binding to dEsyt in its ability to modulate MCS number and distance in the context of an *in vivo* signalling system. Since the protein has been suggested to play dual roles at the contact interface *in vitro*—that of a tether and a lipid transfer protein (Idevall-Hagren et al., 2015; Saheki et al., 2016), our findings emphasise the possibility that Ca^2+^ binding to this protein may modulate such function, in particular the tethering role at the contact interface in an *in vivo* system. While further studies are required to elucidate the *in vivo* significance of these calcium binding residues in maintaining the protein’s lipid transfer activity, overall our study demonstrates the modulation of MCS density by the output of an ongoing signal transduction pathway in cells.

## Acknowledgements

This work was supported by the Department of Atomic Energy, Government of India, under Project Identification No. RTI 4006, a Wellcome-DBT India Alliance Senior Fellowship to PR (IA/S/14/2/501540) and a Wellcome-DBT India Alliance Early Career Fellowship to SM (IA/E/17/1/503653). VRN was supported by a fellowship from the Indian Council for Medical Research (2021-9971/CMB-BMS). We thank the NCBS Imaging Facility, Electron microscopy and Drosophila facilities for support.

## Materials and Methods

### Fly stocks

In a constant temperature laboratory incubator, flies (*Drosophila melanogaster*) were reared on rich medium (corn flour, black jaggery, agar, dry yeast, propionic acid, methyl parahydroxy benzoate, orthophosphoric acid) at 25°C and 50% relative humidity. Except for the few pulses of light that the flies were exposed to when the incubator doors were opened, the environment inside the incubator was completely dark. The experiments were conducted under two different lighting conditions: constant illumination and constant darkness. Incubators were kept at 25°C with a white light source (light intensity: 5000 lux) throughout the duration of the experiments involving continuous illumination. Gal4-UAS system was used for the selective activation of the transgene spatially and temporally for targeted gene expression.

### Sequence alignment

The C2 domains of dEsyt (CG6643), human synaptotagmin (EAW97344.1), human extended-synaptotagmin 1 (NP_001171725.1) and human extended-synaptotagmin 2 (NP_065779.1) were aligned using clustalO. The conservation scores on the alignment were mapped using Jalview.

### Model of C2A domain of dEsyt protein

3-dimensional structural model of C2A domain of dEsyt protein was obtained using homology modelling method with X-ray crystal structure of human Esyt2 C2A domain (PDB ID: 4npk) as template. 20 models were obtained using MODELLER. The model with the least energy was selected for further analysis. The residues in dEsyt C2A domain that interact with calcium were analysed using PYMOL. MIB, metal ion binding site prediction tool was used to check the affinity and metal ion coordination for calcium ions with dEsyt C2A domain.

### Optical neutralisation

The flies reared under experimental conditions were immobilised by cooling on ice, carefully decapitated and fixed on the microscope slide using a drop of colourless nail varnish. A drop of immersion oil was used to neutralise the cornea’s refractive index, which was then viewed with a 40X oil immersion objective. CellSens software was used for digital image acquisition and documentation.

### Scoring retinal degeneration

A total of 50 ommatidia from 5 distinct flies of each genotype were analysed for each time point in order to produce a quantitative indicator of degeneration. The single, central UV sensitive photoreceptor that didn’t show any light dependent retinal degeneration was used as the reference and the rest of the photoreceptors were scored. Each rhabdomere that appeared to be wild type received a score of one. As a result, the control photoreceptors will receive a score of 7, whereas the mutant photoreceptors that are degenerating will receive a score ranging from 1 to 7. GraphPad Prism software was used to analyse the data and plot the graph.

### Immunohistochemistry

Retinae were dissected in PBS and fixed for 30 minutes at room temperature with 4% paraformaldehyde in PBS with 1 mg/ml saponin. After fixation, the samples were rinsed thrice in PBS with 0.3% Triton X-100 (0.3% PBTX), then incubated for 2 hours at room temperature with blocking solution (5% Fetal Bovine solution in PBTX). The samples were incubated overnight with the appropriate primary antibody [rat anti-RDGB (lab generated); 1:300, mouse anti-GFP (Santa Cruz, sc-9996); 1:300, rabbit anti-mCherry (Thermo Fisher Scientific-PA5- 34974); 1:300] at 4°C. The samples were washed three times with 0.3% PBTX and incubated for 4 hours at room temperature with appropriate secondary antibodies [Molecular Probes’ Alexa Fluor 568 anti-rabbit (A10042); 1:300, Molecular Probes’ Alexa Fluor 633 anti-rat (A21094); 1:300, Molecular Probes’ Alexa Fluor 488 anti-mouse (A11001); 1:300]. Along with the secondary antibody incubation, Alexa Fluor 488–Phalloidin (Invitrogen, A12379; 1:200) and Alexa Fluor 568–Phalloidin (Invitrogen, A12380; 1:200) was used to mark F-actin. Samples were washed thrice with 0.3% PBTX, followed by one final wash in PBS and were mounted with 70% glycerol in 1X PBS. The whole-mounted preparations were imaged under 60X 1.4 NA objective, in Olympus FV 3000 microscope.

### Transmission Electron Microscopy

The fly heads of mentioned genotypes were cut and immersed in 2% osmium tetroxide, kept at 4°C for 1 hour followed by incubation at 40°C for 4 days. Specimens were washed with distilled water, stained enbloc with uranyl acetate (0.5% in distilled water) for 3 hours. After washing with distilled water, specimens were subjected to dehydration step and embedded in epon. Ultrathin sections of 60 nm were cut and grids were subjected to post staining with 2% uranyl acetate (in 70% ethanol) and Reynold’s lead solution. Sections were imaged at 120 KV on a Tecnai G2 Spirit Bio-TWIN (FEI) electron microscope and at 200 KV on a Talos F200 G2 (FEI) electron microscope.

### Scoring MCS density and Contact site distance

For scoring MCS density/photoreceptor cell total of 25-30 cells were taken to conduct analysis for R1-R6. Using free hand line tool of image J, length of MCS (µm) /the total length of the base of the rhabdomere (µm) were calculated. Fractions of MCS coverage were multiplied with 100 to show the percentage. For contact site distance, using straight line tool of image J, the distance between the base of rhadbomere to ER was manually analysed and quantified. Graphs were plotted using Prism 8 software.

### Cell culture, transfection and Immunofluorescence

Schneider’s insect medium (HiMedia) was used to culture S2R+ cells, which was supplemented with 10% Fetal Bovine Serum and antibiotics Penicillin and Streptomycin. Effectene (Qiagen) was used to transfect cells according to the manufacturer’s instructions. Post 24 h of transfection, cells were fixed with 4% paraformaldehyde (Electron Microscopy Sciences) and imaged to observe for GFP fluorescence using a 60X 1.4 NA objective, in Olympus FV 3000 microscope.

### Statistical analysis

Unpaired two tailed t test or ANOVA, followed by Tukey’s multiple comparison test, were carried out where applicable.

